# Effect of prophylactic use of intra-nasal oil formulations in the hamster model of Covid-19

**DOI:** 10.1101/2021.06.25.449990

**Authors:** Zaigham Abbas Rizvi, Manas Ranjan Tripathy, Nishant Sharma, Sandeep Goswami, N Srikanth, J L N Sastry, Shailendra Mani, Milan Surjit, Amit Awasthi, Madhu Dikshit

## Abstract

Severe acute respiratory syndrome coronavirus 2 (SARS-CoV2) infection initiates with viral entry in upper respiratory tract leading to coronavirus disease 2019 (Covid-19). Severe Covid-19 is characterized by pulmonary pathologies associated with respiratory failure. Thus, therapeutics aimed at inhibiting entry of the virus or its internalization in the upper respiratory tract, are of interest. Herein, we report the prophylactic application of two intra-nasal formulations provided by the National Medicinal Plant Board (NMPB), Anu oil and Til tailya in SARS-CoV2 infection hamster model. Prophylactic nasal instillation of these oil formulations exhibited reduced viral load in lungs, and resulted in reduced body weight loss and pneumonitis. In line with reduced viral load, histopathlogical analysis revealed a reduction in lung pathology in Anu oil group as compared to the control infected group. However, Til tailya group did not show a significant reduction in lung pathology. Furthermore, molecular analysis using mRNA expression profiling indicated reduced expression of pro-inflammatory cytokines genes, including Th1 and Th17 cytokines for both the intra-nasal formulations as a result of decreased viral load. Together, the prophylactic intra-nasal application of Annu oil seems to be useful in limiting both the viral load and disease severity disease in SARS-CoV2 infection in hamster model.

## Introduction

Since the first report from Wuhan in December 2019, a number of COVID-19 incidences have exploded around the globe and were declared a pandemic by WHO ^1,2^ (https://www.ecdc.europa.eu/en/geographical-distribution-2019-ncov-cases). As of 26^th^ May 2021, the total number of coronavirus infection incidences was 168,701,661 with around 3,502,901 deaths globally with 312,146 mortalities in India alone. The majority of the coronavirus cases are asymptomatic and do not require aggressive treatment. However, an estimated 13.8 % of the infected individuals are at risk of developing a severe form of COVID-19 which could be characterized by either one or all of the following COVID-19 symptoms viz respiratory distress, high fever, loss of taste and smell, and diarrhea^1,2^. In addition, up to around 6% of COVID-19 cases end up with respiratory failure due to cytokine storm, cardio-vascular complications and multiple organ failure (https://www.who.int/emergencies/diseases/novel-coronavirus-2019) ^2,3^. As is known for other respiratory viruses, SARS-CoV2 initially infects the upper respiratory tract and then rapidly spreads to the lower respiratory tract^1^. During an active infection, the virus can be transmitted and spread from both symptomatic as well as asymptomatic individuals via respiratory droplets generated through coughing, sneezing, or hyperventilation via the airborne route^3,4^.

Global health research has primarily focused on vaccine development against COVID-19 with active vaccination being the current strategy to protect COVID-19 related mortalities^**5**,**6**^. Given the emergence of new SARS-CoV2 variants, the protective efficacy of vaccines could be reduced, hence therapeutics that may prevent viral entry, replication and transmission are highly desirable. In line with this, pharmacological agents such as intra-nasal delivery of TLR2/6 agonist, or lipopeptide agents, intra-nasal administration of neutralizing antibodies and intra-nasal gene therapy are currently being explored as potential strategies to inhibit the host-pathogen interaction and limit the infection^**7–12**^. Intra-nasal corticosteroid spray for the recovery of the sense of smell is in clinical trials for COVID-19 patients^**13**^. Since pharmaceutical drugs may have many off-target effects, therapeutics based on herbal extracts have recently gained much attention^**14–17**^. Here, we evaluated the efficacy of two Ayurvedic intra-nasal (herbal) oil formulations viz Anu oil and Til Tailya^**18**^, in hamsters SARS-CoV2 challenge model. Hamsters are the preferred animal model to study SARS-CoV2 infection as they mimic infection of the upper and lower respiratory tract similar to humans and develop SARS-CoV2 related pathologies^**19–21**^.

Sesame oil (Til taila, TT) is oil derived from a plant (*Sesamum indicum*), is a classical ayurvedic medicine mentioned in Charak Samhita (https://niimh.nic.in/ebooks/ecaraka/). On the other hand, a classical Ayurvedic medicine, Anu Tailya was used by Maharishi Charak more than 5000 years ago for therapeutic purposes. Anu taila consists of oil derived from several important medicinal plants like; Nāgarmothā (*Cyperus scariosus*), Jīvantī (*Leptadenia reticulata*), Śweta candana (*Santalum album*), Jala (*Pavonia odorata*), Pṛśniparṇī (*Uraria picta*), Bela (*Aegle marmelos*), Devdāru (*Cedrus deodara*), Dāruharidrā (*Berberis aristata*), Tejpatra (*Cinnamomum tamala*), Dālacīnī (*Cinnamomum verum*), Kamala keṣara (*Nelumbo nucifera*), Sevya (*Chrysopogon zizanioides*), Viḍañga (*Embelia ribes*), Utpala (*Nymphaeanouchali*), Anantmūla (*Hemidesmus indicus*), Tila Taila (*Sesamum indicum*), Muleṭhī (*Glycyrrhiza glabra*), Plawa (*Cyperus platyphyllus*), Agarū (*Aquilaria agallocha*), Śatāvarī (*Asparagus racemosus*), Bṛhatī (*Solanum indicum*), Kaṇṭakārī (*Solanum surattense*), Surbhi (*Pluchea lanceolata*), Śālaparṇī (*Desmodium gangeticum*), Truṭi (*Elettaria* cardamomum), Reṇukā (*Vitex agnus-castus*), and Ajadugdha (Duraipandi and Selvakumar, 2020; see the enclosed supplement information). Here we report that intra-nasal instillation of both Til tailya and Anu oil limited the viral entry and replication into the lungs associated with SARS-CoV2 infection in hamsters. However, Anu oil but not Til tailya was able to rescue the pneumonitis and lung injury partly due to suppression of inflammatory cytokine response.

## Results

### Prophylactic use of intranasal instillation of Ayush oil formulations prevents SARS-CoV2 infection and associated gross clinical parameters

SARS-CoV2 infection in hamsters peaks between 4-5 days and is characterized by a reduction in body weight and appearance of pneumonitis in the lungs and splenomegaly ^19–22^. These defined gross clinical parameters were recorded between all the groups i.e. uninfected control (UI), infected control (I), and infected hamsters receiving either Til tailya (I + TT) or Anu oil (I + AO) intra-nasal formulations (50µl/ nostril/ day). Treatment of TT and AO was started 5 days before SARS-CoV2 live virus challenge and was continued till the end of the experiment i.e. 4 days post infection (dpi), as presented schematically in the study design (**Fig. 1A)**. In line with the earlier published reports, a decrease in body weight of SARS-CoV2-infected hamsters was observed with 5-8% body weight reduction on 4 days dpi. Hamsters receiving TT or AO, before SARS-CoV2 infection, did not lose body weight as was observed in the SARS-CoV2-infected group (**Fig. 1B & 1C**). SARS-CoV2 infection in hamster model is characterized by lung inflammation, pneumonitis, and cytokines release^1,23,24^. To further understand the other parameters related to lung-associated pathologies, gross morphological changes were compared between healthy, SARS-CoV2 -infected and SARS-CoV2 infected plus oil formulations groups.

**Figure 1.**
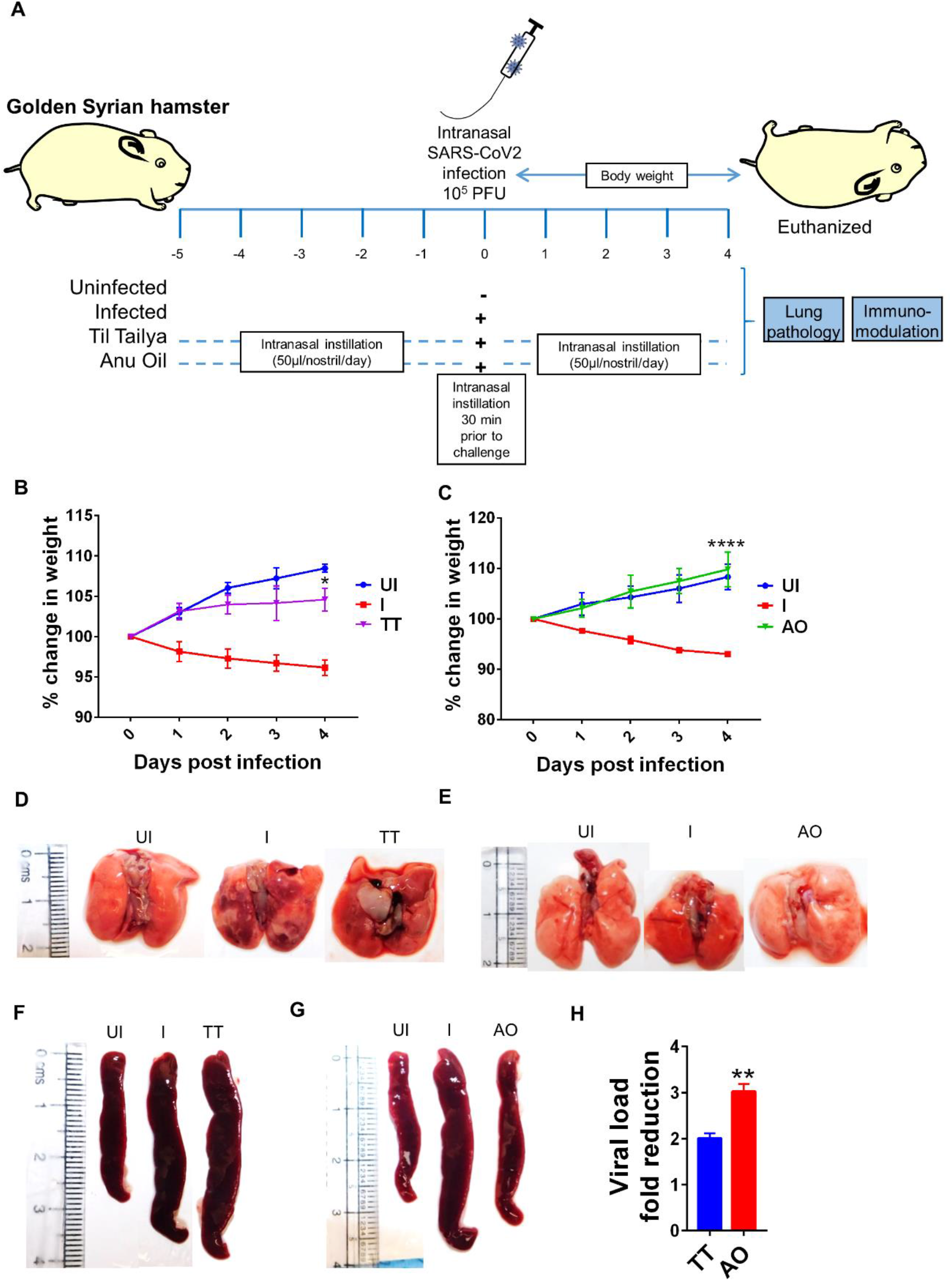
Effect of intra-nasal instillation of Anu oil and Til tailya on gross clinical parameters and lung viral load in SARS-CoV2 infected hamsters. (A) Depicts the schematic outlines the study design. Prophylactic treatment regime was adopted for Anu Oil (AO) and Til tailya (TT) with each animal receiving (50µl/ nostril/ day) intra-nasal instillation of Anu oil, Til Tailya or mock control 5 days before challenge and then continued till after infection till end point (ie 4 days post infection). One group was challenge and received mock control (I); the other group of animals were unchallenged healthy control (UI). On the day of challenge, the animals were given intra-nasal instillation 30 min prior to challenge with SARS-CoV2. (B & C) Line graph showing mean % change in body weight post infection ± standard error mean (SEM). (D & E) Images of the excised lungs showing gross morphology with pneumonitis region (dark red patches). (F & G) Images of excised spleen indicating changes in the spleen length (H) Bar graph showing mean fold reduction in lung viral load ± SEM as compared to infected (I) control. **P < 0.01 (t-test).

Results obtained indicated a reduction in the regions of pneumonitis in the excised lungs of AO group, but not TT group, as compared to infection control (**Fig. 1D & 1E**). As reported earlier, splenomegaly is one of the critical parameters indicative of active infection^20^. Thus we tested the splenomegaly between different groups and found that AO, but not TT, showed inhibition in splenomegaly as compared to the SARS-CoV2 infected hamsters (**Fig. 1F and 1G)**. We also evaluated the lung viral load at 4 dpi and calculated the fold reduction in viral load in AO and TT treated groups as compared to the SARS-CoV2 infected groups. Our data indicate that compared to the SARS-CoV2 infected group, viral loads in AO and TT treated groups were ∼3 and ∼2 fold less respectively (**Fig. 1H)**. Together, these data indicated that prophylactic use of intra-nasal instillation of TT and AO resulted in decreased lung viral load with AO group showing better protection in gross clinical parameters.

### Prophylactic use of Anu Oil reduces SARS-CoV2-induced lung pathology in hamsters

Since gross clinical parameter viral load data suggested protection from SARS-CoV2 infection in AO and TT groups, we set out to study the mitigation of pulmonary pathologies such as lung injury, alveolar epithelial injury, bronchitis, pneumonitis and inflammation using histological analysis^20,25–27^. Haematoxylin and eosin (H & E) stained lung data showed a reduction in alveolar epithelial injury, inflammation and pneumonitis in SARS-CoV2 infected-AO treated hamsters as compared to SARS-CoV2 infected hamsters. There was no sign of bronchitis in SARS-CoV2 infected-AO treated hamsters with overall significant mitigation in disease score as compared to SARS-CoV2 infected hamsters (**Fig. 2A and 2B)**. Hamsters treated with TT, however, showed little or no improvement in lung injury and overall disease score as compared to the infected control **(Fig. 2A and 2B)**. Together, lung pathology associated with SARS-CoV2 was found to be resolved in AO group but not in TT group with around 1.5 fold rescue in disease index in AO group as compared to the infected control group.

**Figure 2.**
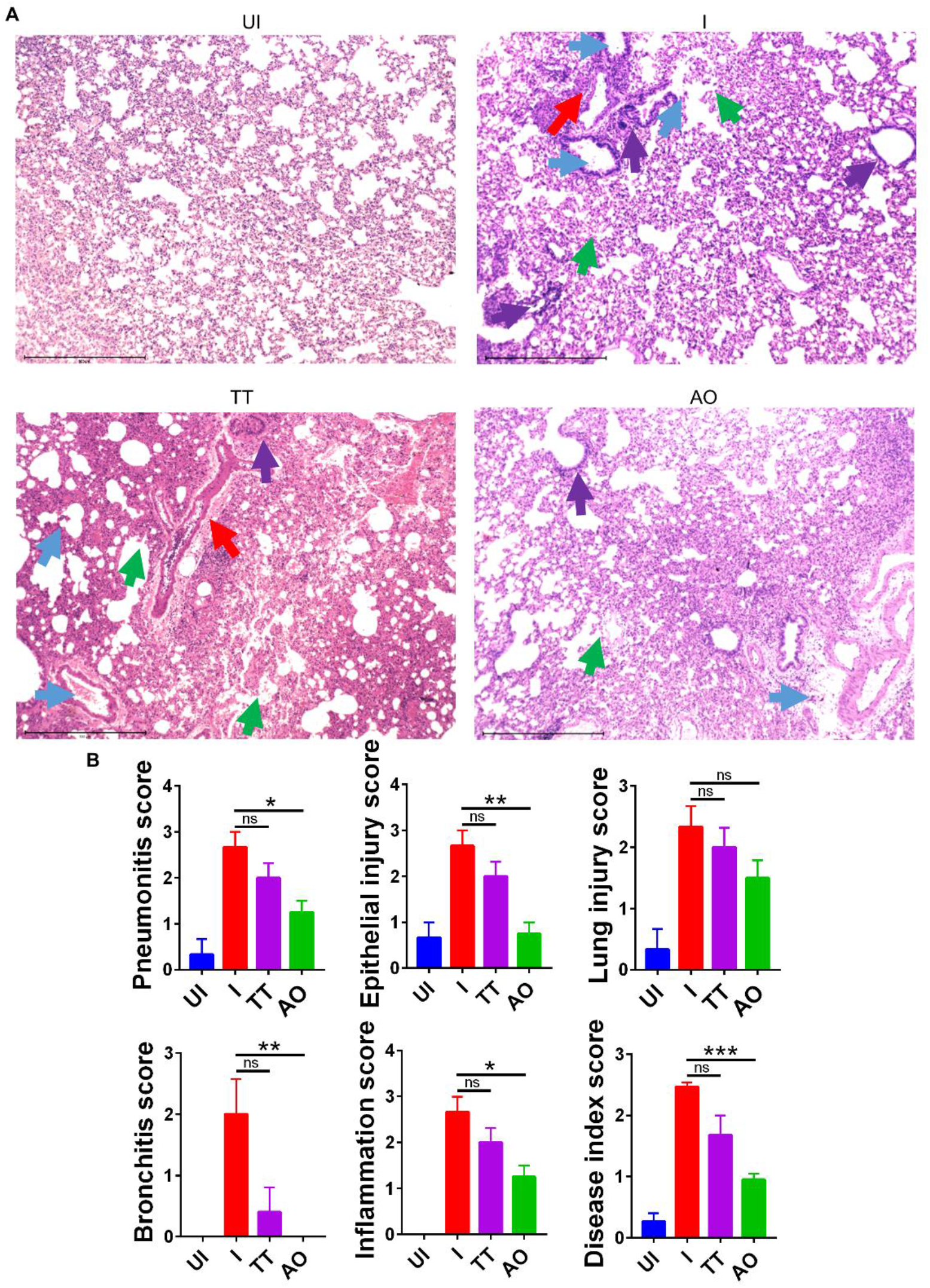
H & E stained lung sections showing histopathology and its assessment. (A) images of H&E stained lungs at 40X magnification showing regions of pneumonitis (blue arrow), bronchitis (red arrow), epithelial injury (green arrow) and inflammation (purple arrow) along with their (B) histological score for pneumonitis, inflammation, lung injury, alveolar epithelial cells, bronchitis and overall disease score for different groups UI, I, TT and AO on day 4 post infection. ^*^P < 0.05, ^**^P < 0.01, ^***^P < 0.001 (one-way anova).

### Prophylactic use of Anu oil prevent lung injury in hamsters associated to SARS-CoV2 infection

Lung histology data indicated overall improved histological changes of SARS-CoV2 infected-AO treated groups as compared to SARS-CoV2 infected group. These data compelled us to explore the mechanism involved in the rescue of lung pathologies. Injury to the pulmonary region is characterized by elevated expression of surfactant D (sftp-D), increased mucus (muc-1) secretion as well as increased expression of eotaxin which promotes infiltration of granulocytes and mast cells and an increased risk of pulmonary thrombosis as seen in COVID-19 patients^28–32^. We observed elevated levels of sftp-D, muc-1, eotaxin, muc-1, chymase, tryptase and plasmonigen activator inhibitor-I (PAI-1: a key factor for lung fibrosis) in the lungs of infected hamsters (**Fig. 3A, 3B and 3C)**. However, prophylactic intra-nasal use of AO in SARS-CoV2-infected hamster significantly reduced the mRNA expression of these lung injury genes as well as genes that are required for chemotaxis of granulocytes and function of mast cells (**Fig. 3A and 3B)**. Prophylactic use of AO in SARS-CoV2 infected hamsters did not reduce PAI-1 expression (**Fig. 3C**). In contrast to AO, TT data showed no reduction in lung injury as compared to infected control (**Fig. 3A)**. Surprisingly, there was a marked increase in mast cell markers in TT group as compared to the infected control (**Fig. 3B)**. Overall, consistent with our lung histology data, a profound reduction in the expression of lung injury genes and mast cell markers as compared to the infected control was observed in AO treated groups.

**Figure 3.**
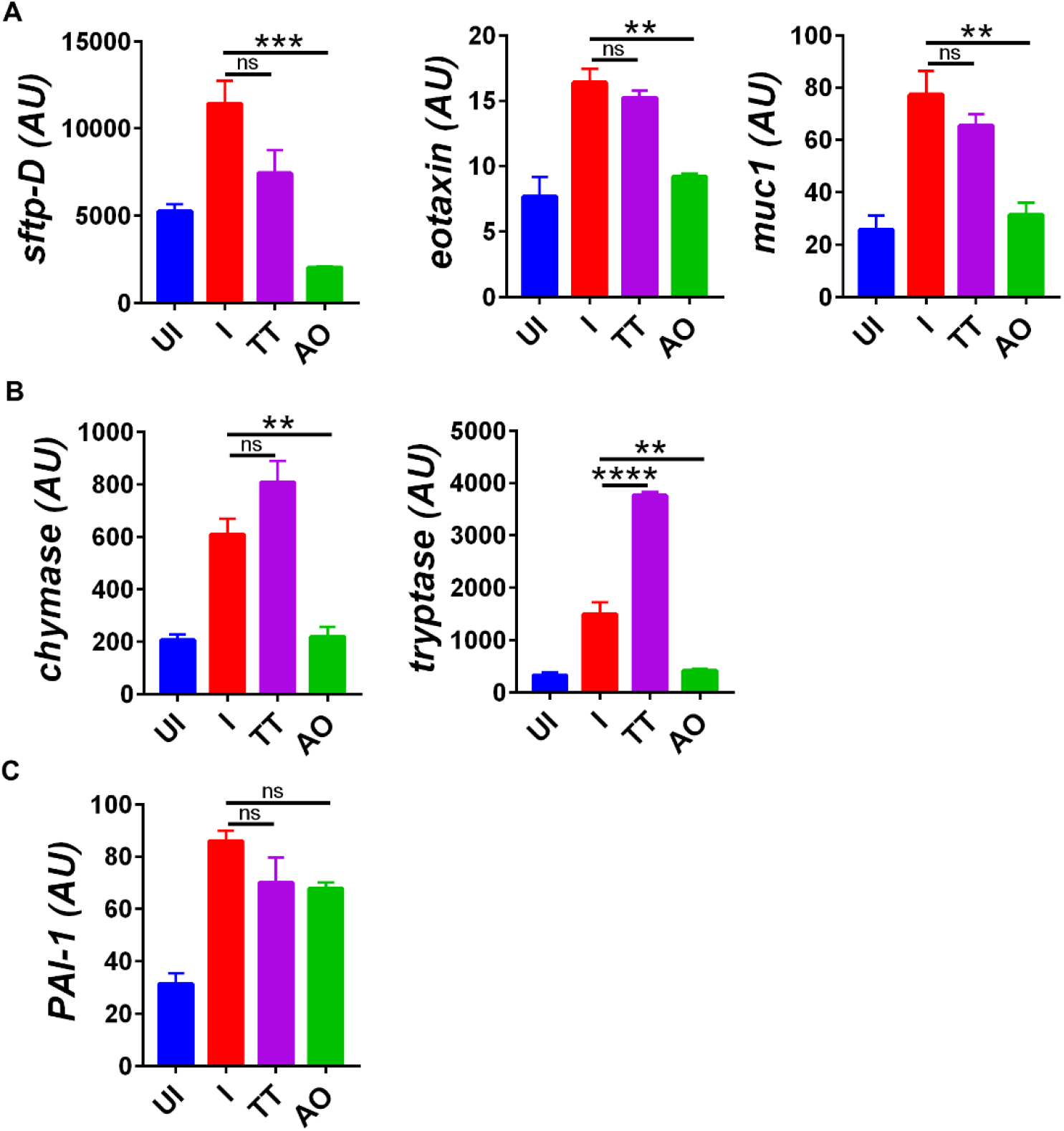
Changes in mRNA expression of genes involved in lung injury upon AO or TT intra-nasal administration in hamsters infected with SARS-CoV2. Relative mRNA expression profiling was carried out in UI, I, TT and AO lung samples for (A) lung injury genes (B) mast cell activation markers (C) thrombosis factor. Mean ± SEM. ^**^P < 0.01, ^***^P < 0.001, ^****^P < 0.0001 (one-way anova).

### AO and TT treatment inhibits the expression of SARS-CoV2-induced pro-inflammatory cytokines

COVID-19 related respiratory distress is associated with inflammation in the lungs. The increase in lung inflammation is characterized by the secretion of pro-inflammatory cytokines in COVID-19 patients^24,33–35^. Cytokine expression data from splenocytes indicates elevated expression of Th1 cytokines (IFNγ and TNFα), Th2 cytokine (IL-4, IL-13), Th17 cytokine (IL-17A) and various others pro-inflammatory cytokine-like IL-6 in SARS-CoV2 -infected as compared to uninfected hamsters (**Fig. 4A, 4B, 4C and 4D)**. Prophylactic intra-nasal installation of AO and TT in SARS-CoV2 infected hamsters resulted in the reduction in Th1, Th17 cell cytokines together with pro-inflammatory cytokines expression (**Fig. 4A, 4B and 4D)**. However, surprisingly only TT but not AO was able to reduce the Th2 cytokines gene expression (**Fig. 4A and 4C)**. Further, we found an elevated IFNy secretion in both AO and TT treated animals (**Fig. 4A and 4C)**. Together, we show that AO and TT reduced the pro-inflammatory cytokines.

**Figure 4.**
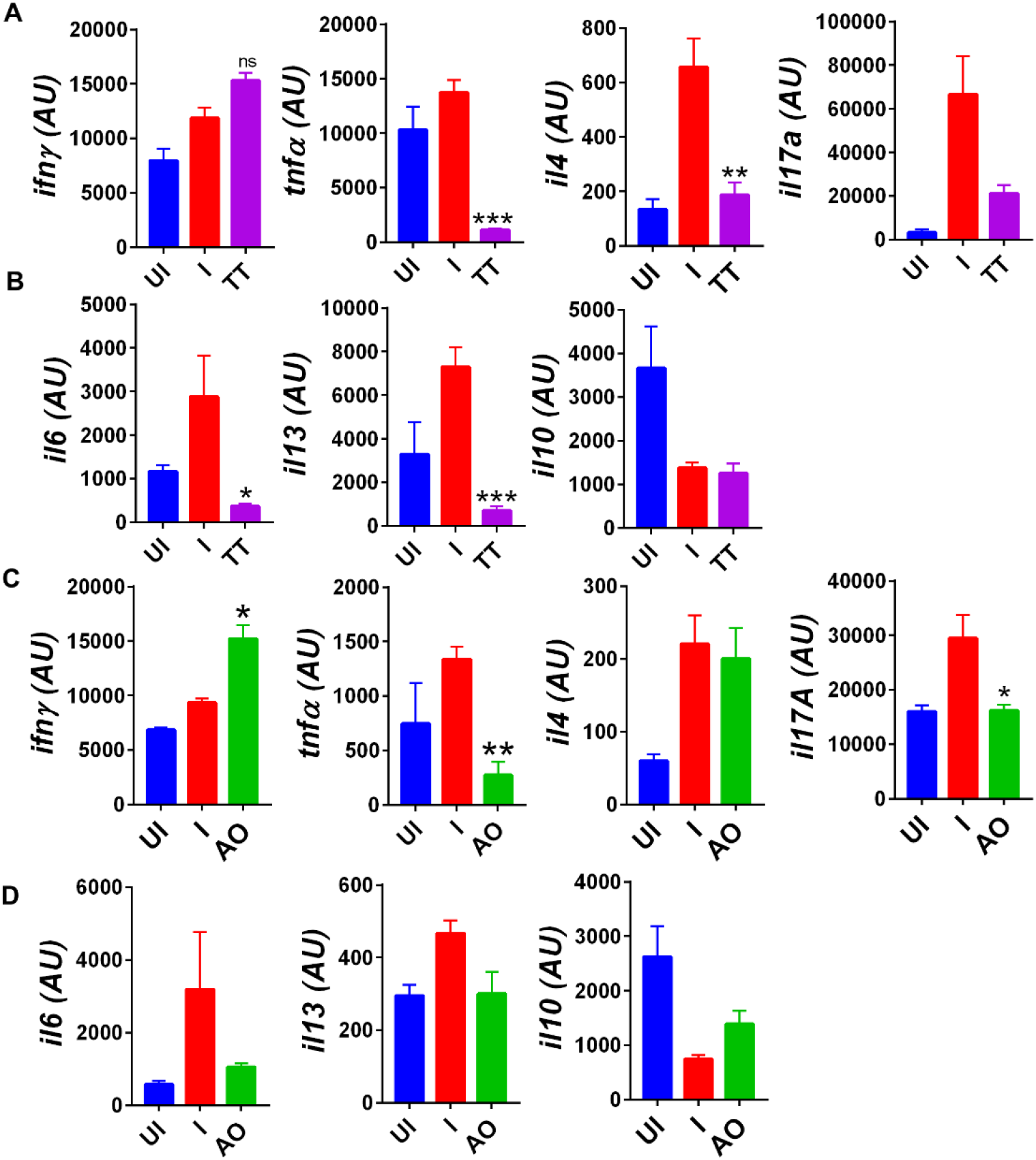
Immuno-modulatory effect of TT and AO on cytokine expression in splenocytes. Relative mRNA expression profiling was carried out in UI, I, TT and AO splenocytes samples for (A) T helper cell cytokines for TT samples (B) pro-inflammatory and anti-inflammatory cytokines for TT samples (C) T helper cell cytokines for AO samples (D) pro-inflammatory and anti-inflammatory cytokines for AO samples. Bar graph showing mean ± SEM. ^*^P < 0.05 ^**^P < 0.01, ^***^P < 0.001, (one-way anova).

### Anu Oil intra-nasal formulation shows more protective efficacy against SARS-CoV2 infection in hamsters as compared to Til Tailya intra-nasal formulation

We summarize the finding of our study and provide the first evidence that intra-nasal formulation such as Anu Oil and Til tailya limits viral entry and replication. However, only Anu Oil but not Til tailya was effective in reducing the SARS-CoV2 associated pulmonary pathologies and lung injuries even though both AO and TT were effective in reducing the pro-inflammatory cytokine response in hamsters (**Fig.5**).

**Figure 5.**
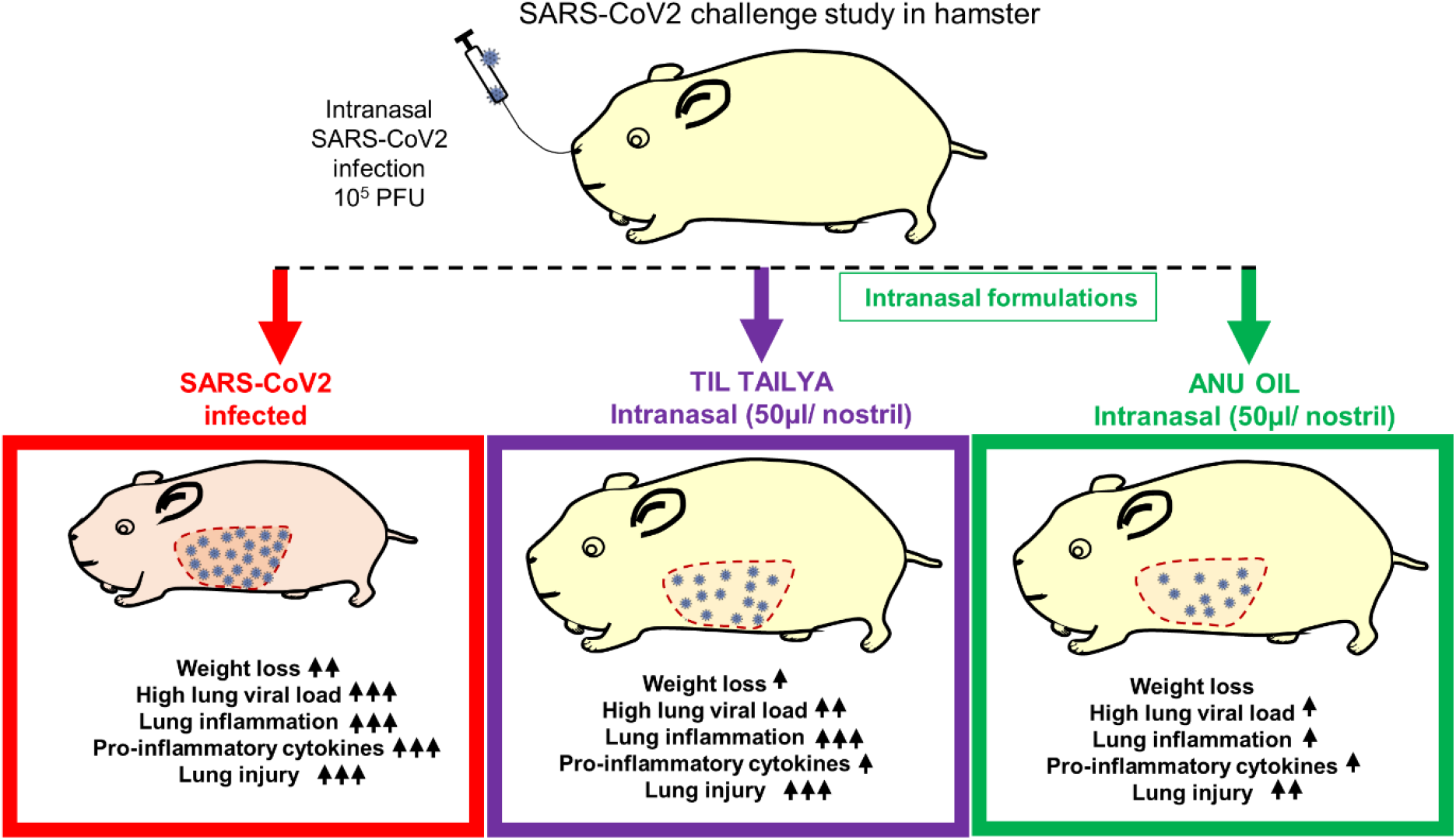
Graphical summary representing the important findings of the study. Diagramatic representation of the effect of pretreatment of intra-nasal Til tailya and Anu Oil against SARS-CoV2 associated pathologies in golden Syrian hamster.

## Discussion

Advances in vaccine development against SARS-CoV2 Wuhan strain and aggressive vaccination strategy have significantly reduced the SARS-CoV2 infection and COVID-19 deaths worldwide ^5,6,36^. However, the recent rise in variant strains that show high transmission and disease severity is of concern, primarily, due to the reduced efficacy of virus neutralization in vaccinated individuals for some variant strains^37–39^. Therefore, therapeutic options are paramount in combating COVID-19.

Though several repurposed drugs are currently being used for the treatment of COVID-19 patients but have limited efficacy. Herbal medicines or medicines derived from herbal extracts offer a safer alternative therapy due to their prolonged human use, acceptability with lesser side effects^15–17^. Recently, Chinese traditional medicine has gained popularity due to its anti-viral efficacy in in-vitro and animal models of SARS-CoV2 ^15,40–43^. In India, going back to more than 3000 years, Ayurvedic medicines are considered useful for both lifestyle disorders and infectious conditions. The word Ayurveda is derived from two Sanskrit words ayur (life) and veda (knowledge) ^44–49^.

In the current study, we describe the efficacy of prophylactic use of two intra-nasal Ayurvedic oil formulations using hamster model for SARS-CoV2 challenge. Hamsters are one of the best small animal models for SARS-CoV2 infection which mimic the viral entry, replication of the upper and lower respiratory tract of humans^20,21^. Hamsters receiving Anu Oil or Til tailya intranasally before SARS-CoV2 infection showed reduced weight loss. It has been shown earlier that even low dose of SARS-CoV2 intra-nasal infection in hamster could result in pneumonitis and lung pathologies^20^. Therefore, it might be possible that even though both Til tailya and Anu Oil can limit SARS-CoV2 entry and replication by some degree but only Anu oil group showed a greater reduction in the lung viral load (around 3 folds). Lung histology data corroborated with the gross parameters findings showing lesser SARS-CoV2 related histopathology in Anu Oil treated hamsters while there was little or no protection in Til Tailya group. Pulmonary damage and pneumonitis have been reported as the major cause of respiratory failure in patients suffering from a severe form of SARS-CoV2 infection ^24^. Prophylactic use of Anu Oil showed significant reduction in the overall disease index as compared to the infected control, suggesting some degree of protection against SARS-CoV2 induced lung injury and pathology. Since hamster antibodies are not available commercially we carried out mRNA expression profiling. Our, mRNA expression data for lung injury genes suggest that Anu oil intervention also reduced the expression of lung injury markers and significantly reduced the lung inflammation, indicating that Anu Oil was able to protect against the pulmonary damage caused by SARS-CoV2 infection. Finally, we studied the expression of cytokines to understand if intra-nasal formulation could help prevent the inflammatory cytokine response within the lung. Interestingly, both Anu Oil and Til Tailya were able to limit the expression of pro-inflammatory cytokines as compared to the infected hamsters.

Taken together, in the current study we report using hamster SARS-CoV2 model that prophylactic intra-nasal treatment with Anu Oil as well as Til Tailya reduced the lung viral load. However, SARS-CoV2 related pulmonary pathologies were prevented only in Anu Oil-treated hamsters as demonstrated by histopathological lung injury scores and expression of injury markers and inflammatory cytokines. The protection against SARS-CoV2 infection and related pathologies seem to be in part due to the significant reduction in the viral entry and replication in the upper respiratory tract. This preclinical study in the hamster model points to the prophylactic potential of intra-nasal Annu oil in Covid and necessitates further studies to understand its observed effect.

## Methods

Seasame oil and Anu Taila (a poly herbal medicine) used in the study were prepared as per pharmacopial standards and were provided by National Medicinal Plant Board for the study.

### Animal Ethics and biosafety statement

6-8 weeks old female golden Syrian hamsters were acclimatized in biosafety level-2 (BSL-2) for one week and then infected in Animal BSL3 (ABSL-3) institutional facility. The animals were maintained under 12 h light and dark cycle and fed a standard pellet diet and water ad libitum. All the experimental protocols involving the handling of virus culture and animal infection were approved by RCGM, institutional biosafety, and IAEC animal ethics committee (IAEC/THSTI/105).

### Virus culture and titration

SARS-Related Coronavirus 2, Isolate USA-WA1/2020 virus was grown and titrated in Vero E6 cell line cultured in Dulbecco’s Modified Eagle Medium (DMEM) complete media containing 4.5 g/L D-glucose, 100,000 U/L Penicillin-Streptomycin, 100 mg/L sodium pyruvate, 25mM HEPES, and 2% FBS. The stocks of the virus were plaque purified at THSTI IDRF facility inside ABSL3 following institutional biosafety guidelines.

### SARS-CoV2 infection in golden Syrian hamster and Ayush herbal extracts dosing regime

6-9 weeks golden Syrian hamsters were procured from CDRI and quarantined for one week at small animal facility (SAF), THST before starting the experiment. One group received intra-nasal installation of Anu oil while the other group received intra-nasal installation of Til Tailya (50ul/ nostril/ day) started 5 days before infection and continued till 4 days post-infection (DPI). On the day of the challenge, intra-nasal administration of Anu oil and Til Tailya was carried out 30 min before infection. On the day of the challenge, the animals were shifted to ABSL3. Intra-nasal infection with live SARS-CoV2 10^5^PFU/ 100μl or with DMEM mock control was established with the help of catheter under mild anesthetized by using ketamine (150mg/kg) and xylazine (10mg/kg) intraperitoneal injection inside ABSL3 facility. All the experimental protocols involving the handling of virus culture and animal infection were approved by RCGM, institutional biosafety and IAEC animal ethics committee.

### Gross clinical parameters of SARS-CoV2 infection

All infected animals were euthanized on 4 days post infection at ABSL3. Changes in body weight were observed on each day post-challenge and plotted as percent change in the body weight. Post sacrifice, lungs and spleen of the animals were excised and imaged for gross morphological changes. Left lower lobe of the lung was fixed in 10% formalin and used for histological analysis. The remaining part of the lung left lobe was homogenized in 2ml Trizol solution for viral load estimation. Spleen was homogenized in 2ml of Trizol solution. The trizol samples were stored immediately at -80 °C until further use. Blood of the animals was drawn through direct heart puncture and serum was isolated and stored at -80 °C until further use.

### Viral load

For viral load, estimation lungs were homogenized in Trizol reagent (Invitrogen) and their supernatant was collected after centrifugation at 4000 rpm for 15 min at 4 °C. Thereafter, RNA was isolated by Trizol-Choloform method and RNA yield was quantitated by nano-drop. 1 µg of total RNA was then reverse-transcribed to cDNA using the iScript cDNA synthesis kit (Biorad; #1708891) (Roche). Diluted cDNAs (1:5) were used for qPCR by using KAPA SYBR® FAST qPCR Master Mix (5X) Universal Kit (KK4600) on Fast 7500 Dx real-time PCR system (Applied Biosystems) and the results were analyzed with SDS2.1 software. Briefly, 200 ng of RNA was used as a template for reverse transcription-polymerase chain reaction (RT-PCR). The CDC-approved commercial kit was used for of SARS-CoV-2 N gene: 5′-GACCCCAAAATCAGCGAAAT-3′ (Forward), 5′-TCTGGTTACTGCCAGTTGAATCTG-3′ (Reverse). Hypoxanthine-guanine phosphoribosyltransferase (HGPRT) gene was used as an endogenous control for normalization through quantitative RT-PCR. The relative expression of each gene was expressed as fold change and was calculated by subtracting the cycling threshold (Ct) value of hypoxanthine-guanine phosphoribosyltransferase (HGPRT-endogenous control gene) from the Ct value of the target gene (ΔCT). Fold change was then calculated according to the formula POWER(2,-ΔCT)*10,000 ^50^.

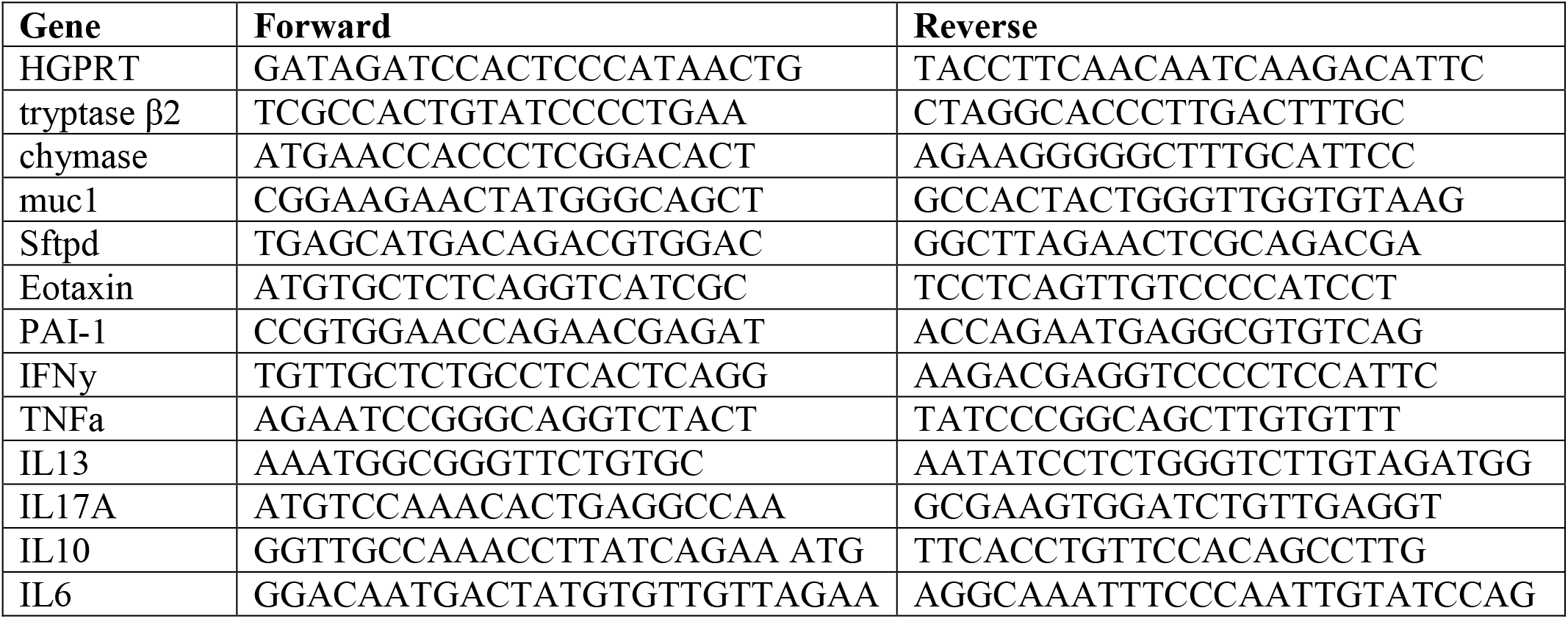

### qPCR

RNA from spleen samples was isolated as described above for the lung samples and cDNA was prepared. Thereafter, the relative expression of each gene was expressed as fold change and was calculated by subtracting the cycling threshold (Ct) value of hypoxantine-guanine phosphoribosyltransferase (HGPRT-endogenous control gene) from the Ct value of target gene (ΔCT). Fold change was then calculated according to the formula POWER(2,-ΔCT)*10,000^50^. The list of the primers is provided as follows.

### Histology

The lung of the euthanized animals was fixed in 10% formalin solution and then embedded in paraffin. Samples embedded paraffin blocks were then cut into 3 µm fine sections and then mounted on silane-coated glass slides. The slides were then stained with hematoxylin and eosin dye as previously described^51^. Each stained sample was then analyzed and captured at 40X magnification. Assessment for the histological score was carried out through blind scoring for each sample by a professional histologist.

### Statistical analysis

Results from the experiments were analyzed and plotted by using Graph pad prism 7.0 software. Graph for percent change in body weight, gene expression, lung histology scores were compared and analyzed by using student t-test or one-way ANOVA with n=5 samples per group. P-value of less than 0.05 and was considered as statistically significant.

## Acknowledgments

MD and AA received financial support for this study from Ayush-DBT (BT/PR40378/TRM/120/486/2020 & NMPB/IFD/GIA/NR/PL/2020-21/53). We acknowledge the financial support from THSTI Core to establish the hamster model for SARS-CoV2 infection. We greatly acknowledge the support and critical inputs of Dr. Pramod Kumar Garg in the manuscript. We acknowledge IDRF (THSTI) for the support at ABSL3 facility. Pankaj Lawania and Prabhanjan Diwedi for their support in conducting ABSL3 animal experiments and Jitender Chandel for providing technical help. Small animal facility and Immunology Core for providing support in experimentation. Hamsters were procured from CDRI, Lucknow. ILBS for support in histological analysis and assessment. The following reagent was deposited by the Centers for Disease Control and Prevention and obtained through BEI Resources, NIAID, NIH: SARS Related Coronavirus 2, Isolate USA-WA1/2020, NR-52281.

## Author Contributions

Conceived, designed and supervised the study: MD, AA; Performed the experiments: ZAR, NS,; ABSL3 procedures: ZAR, MRT, NS; qPCR primer designing and analysis: ZAR; Histology and analysis: ZAR, MRT; Viral load: ZAR, MRT, SG; qPCR: ZAR, MRT, SG; Virus stock: SM; Contributed reagents/materials/analysis tools: AA; Wrote the manuscript: ZAR; Revised the manuscript: MD, AA.

## References

1. Chen, Y. & Li, L. SARS-CoV-2: virus dynamics and host response. The Lancet Infectious Diseases 20, 515–516 (2020).

2. Wang, T. et al. Comorbidities and multi-organ injuries in the treatment of COVID-19. The Lancet 395, e52 (2020).

3. Guan, W.-J. et al. Clinical Characteristics of Coronavirus Disease 2019 in China. N Engl J Med 382, 1708–1720 (2020).

4. Gandhi, R. T., Lynch, J. B. & del Rio, C. Mild or Moderate Covid-19. New England Journal of Medicine 0, null (2020).

5. Dong, Y. et al. A systematic review of SARS-CoV-2 vaccine candidates. Signal Transduction and Targeted Therapy 5, 1–14 (2020).

6. Poland, G. A., Ovsyannikova, I. G., Crooke, S. N. & Kennedy, R. B. SARS-CoV-2 Vaccine Development: Current Status. Mayo Clinic Proceedings 95, 2172–2188 (2020).

7. Boiardi, F. & Stebbing, J. Reducing transmission of SARS-CoV-2 with intranasal prophylaxis. EBioMedicine 63, (2021).

8. Hassan, A. O. et al. A Single-Dose Intranasal ChAd Vaccine Protects Upper and Lower Respiratory Tracts against SARS-CoV-2. Cell 183, 169-184.e13 (2020).

9. Ku, M.-W. et al. Intranasal vaccination with a lentiviral vector protects against SARS-CoV-2 in preclinical animal models. Cell Host & Microbe 29, 236-249.e6 (2021).

10. Kunzelmann, K. Getting hands on a drug for Covid-19: Inhaled and Intranasal Niclosamide. The Lancet Regional Health – Europe 4, (2021).

11. Proud, P. C. et al. Prophylactic intranasal administration of a TLR2/6 agonist reduces upper respiratory tract viral shedding in a SARS-CoV-2 challenge ferret model. EBioMedicine 63, 103153 (2021).

12. Vries, R. D. de et al. Intranasal fusion inhibitory lipopeptide prevents direct-contact SARS-CoV-2 transmission in ferrets. Science 371, 1379–1382 (2021).

13. Abdelalim, A. A., Mohamady, A. A., Elsayed, R. A., Elawady, M. A. & Ghallab, A. F. Corticosteroid nasal spray for recovery of smell sensation in COVID-19 patients: A randomized controlled trial. American Journal of Otolaryngology 42, 102884 (2021).

14. De Pellegrin, M. L. et al. The potential of herbal extracts to inhibit SARS-CoV-2: a pilot study. Clin Phytosci 7, 29 (2021).

15. Jan, J.-T. et al. Identification of existing pharmaceuticals and herbal medicines as inhibitors of SARS-CoV-2 infection. PNAS 118, (2021).

16. Li, S. et al. Edible and Herbal Plants for the Prevention and Management of COVID-19. Front. Pharmacol. 12, (2021).

17. Matveeva, T., Khafizova, G. & Sokornova, S. In Search of Herbal Anti-SARS-Cov2 Compounds. Front. Plant Sci. 11, (2020).

18. Duraipandi, S. & Selvakumar, V. Reinventing nano drug delivery systems for hydrophilic active ingredients in Ayurvedic lipid based formulations containing poly herbal decoction. Journal of Ayurveda and Integrative Medicine 11, 224–227 (2020).

19. Chan, J. F.-W. et al. Simulation of the clinical and pathological manifestations of Coronavirus Disease 2019 (COVID-19) in golden Syrian hamster model: implications for disease pathogenesis and transmissibility. Clin Infect Dis doi:10.1093/cid/ciaa325.

20. Rizvi, Z. A. et al. Immunological and cardio-vascular pathologies associated with SARS-CoV-2 infection in golden syrian hamster. bioRxiv 2021.01.11.426080 (2021) doi:10.1101/2021.01.11.426080.

21. Sia, S. F. et al. Pathogenesis and transmission of SARS-CoV-2 in golden hamsters. Nature 583, 834–838 (2020).

22. Imai, M. et al. Syrian hamsters as a small animal model for SARS-CoV-2 infection and countermeasure development. Proceedings of the National Academy of Sciences 202009799 (2020) doi:10.1073/pnas.2009799117.

23. Afrin, L. B., Weinstock, L. B. & Molderings, G. J. Covid-19 hyperinflammation and post-Covid-19 illness may be rooted in mast cell activation syndrome. International Journal of Infectious Diseases 100, 327–332 (2020).

24. Moore, J. B. & June, C. H. Cytokine release syndrome in severe COVID-19. Science 368, 473–474 (2020).

25. Bao, L. et al. The pathogenicity of SARS-CoV-2 in hACE2 transgenic mice. Nature 583, 830–833 (2020).

26. Lee, A. C.-Y. et al. Oral SARS-CoV-2 Inoculation Establishes Subclinical Respiratory Infection with Virus Shedding in Golden Syrian Hamsters. Cell Reports Medicine 1, 100121 (2020).

27. Leng, L. et al. Pathological features of COVID-19-associated lung injury: a preliminary proteomics report based on clinical samples. Signal Transduction and Targeted Therapy 5, 1–9 (2020).

28. Chatterjee, M., Putten, J. P. M. van & Strijbis, K. Defensive Properties of Mucin Glycoproteins during Respiratory Infections—Relevance for SARS-CoV-2. mBio 11, (2020).

29. Crouch, E. C. Surfactant protein-D and pulmonary host defense. Respiratory Research 1, 6 (2000).

30. Guo, R. F. et al. Regulatory effects of eotaxin on acute lung inflammatory injury. J Immunol 166, 5208–5218 (2001).

31. Prabhakaran, P. et al. Elevated levels of plasminogen activator inhibitor-1 in pulmonary edema fluid are associated with mortality in acute lung injury. Am J Physiol Lung Cell Mol Physiol 285, L20–28 (2003).

32. Xie, G. et al. Hypoxia-induced angiotensin II by the lactate-chymase-dependent mechanism mediates radioresistance of hypoxic tumor cells. Scientific Reports 7, 42396 (2017).

33. Iwasaki, A. & Yang, Y. The potential danger of suboptimal antibody responses in COVID-19. Nature Reviews Immunology 1–3 (2020) doi:10.1038/s41577-020-0321-6.

34. Mathew, D. et al. Deep immune profiling of COVID-19 patients reveals distinct immunotypes with therapeutic implications. Science 369, (2020).

35. Verity, R. et al. Estimates of the severity of coronavirus disease 2019: a model-based analysis. Lancet Infect Dis 20, 669–677 (2020).

36. Poland, G. A., Ovsyannikova, I. G. & Kennedy, R. B. SARS-CoV-2 immunity: review and applications to phase 3 vaccine candidates. The Lancet 396, 1595–1606 (2020).

37. Garcia-Beltran, W. F. et al. Multiple SARS-CoV-2 variants escape neutralization by vaccine-induced humoral immunity. medRxiv (2021) doi:10.1101/2021.02.14.21251704.

38. Planas, D. et al. Sensitivity of infectious SARS-CoV-2 B.1.1.7 and B.1.351 variants to neutralizing antibodies. Nature Medicine 27, 917–924 (2021).

39. Supasa, P. et al. Reduced neutralization of SARS-CoV-2 B.1.1.7 variant by convalescent and vaccine sera. Cell 184, 2201-2211.e7 (2021).

40. Lee, D. Y. W., Li, Q. Y., Liu, J. & Efferth, T. Traditional Chinese herbal medicine at the forefront battle against COVID-19: Clinical experience and scientific basis. Phytomedicine 80, 153337 (2021).

41. Ling, C. Traditional Chinese medicine is a resource for drug discovery against 2019 novel coronavirus (SARS-CoV-2). J Integr Med 18, 87–88 (2020).

42. Xiong, X., Wang, P., Su, K., Cho, W. C. & Xing, Y. Chinese herbal medicine for coronavirus disease 2019: A systematic review and meta-analysis. Pharmacological Research 160, 105056 (2020).

43. Yang, Y., Islam, M. S., Wang, J., Li, Y. & Chen, X. Traditional Chinese Medicine in the Treatment of Patients Infected with 2019-New Coronavirus (SARS-CoV-2): A Review and Perspective. Int J Biol Sci 16, 1708–1717 (2020).

44. Banerjee, S. et al. Immunoprotective potential of Ayurvedic herb Kalmegh (Andrographis paniculata) against respiratory viral infections - LC-MS/MS and network pharmacology analysis. Phytochem Anal (2020) doi:10.1002/pca.3011.

45. Girija, P. L. T. & Sivan, N. Ayurvedic treatment of COVID-19/SARS-CoV-2: A case report. Journal of Ayurveda and Integrative Medicine (2020) doi:10.1016/j.jaim.2020.06.001.

46. Golechha, M. Time to realise the true potential of Ayurveda against COVID-19. Brain Behav Immun 87, 130–131 (2020).

47. Joshi, J. A. & Puthiyedath, R. Outcomes of Ayurvedic care in a COVID-19 patient with hypoxia – A case report. Journal of Ayurveda and Integrative Medicine (2020) doi:10.1016/j.jaim.2020.10.006.

48. Rastogi, S., Pandey, D. N. & Singh, R. H. COVID-19 pandemic: A pragmatic plan for ayurveda intervention. J Ayurveda Integr Med (2020) doi:10.1016/j.jaim.2020.04.002.

49. Subrahmanya, N. K., Patel, K. S., Kori, V. K. & Shrikrishna, R. Role of Kasahara Dashemani Vati in Kasa and Vyadhikshamatva in children with special reference to recurrent respiratory tract infections. Ayu 34, 281–287 (2013).

50. Malik, S. et al. Transcription factor Foxo1 is essential for IL-9 induction in T helper cells. Nat Commun 8, 815 (2017).

51. Rizvi, Z. A., Puri, N. & Saxena, R. K. Evidence of CD1d pathway of lipid antigen presentation in mouse primary lung epithelial cells and its up-regulation upon Mycobacterium bovis BCG infection. PLOS ONE 13, e0210116 (2018).

