## Supplementary file for "Effect of prophylactic use of intra-nasal oil formulations in the hamster model of Covid-19"

**Supplement information**

**ANU TAILA FORMULATION AND its method of preparation**

***Anutaila*** is a very ancient Ayurvedic preparation which is commonly used for *Nasya karma* i.e. errhines / nasal administration. It contains more than 25 herbs which are blended together in the form of decoction. This decoction is slowly infused with sesame oil over a long period of time with the help of controlled heating till the desired quality of oil is obtained. This process is repeated 10 times to have effective potency. *Aja ksheer* (Goat Milk) is also used in the last cycle only. Hence it is said that *Anutaila* is having property of *Maha guna* or *Sarvottama guna* i.e., superior to Oils used for *Nasya karma*. It strengthens the neck, shoulder, chest muscles and improves the capacity of sense organs.

*Brihat trayi* sites *Anutaila* in the context of Nasya many times. *Anutaila* is described by Charak Samhita at su. 5 /63-70, Sushrut Samhita in chi. 4 /28 and Ashtanga Hridaya in Su. 20/36-39. Ashtang sangraha also described 2 types of Anutaila in Su.29 /10-11 and in Anandkand it is cited in Amrutikaran vishranti 18/95-103. These Ayurvedic texts explain Anutaila in different contexts. The process of Anutaila preparation is explained in the following **flow chart** and the general contents of Anutaila are mentioned in the Table below.

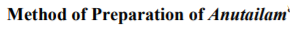

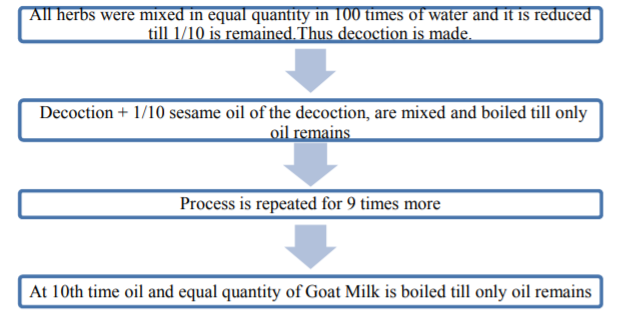

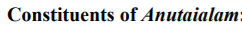

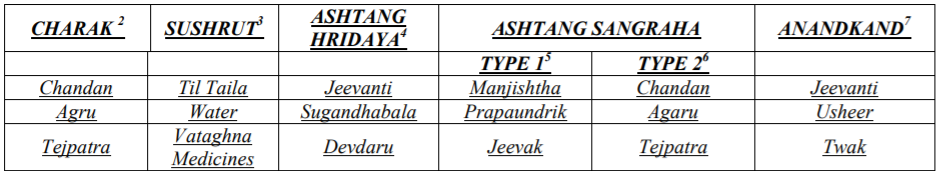

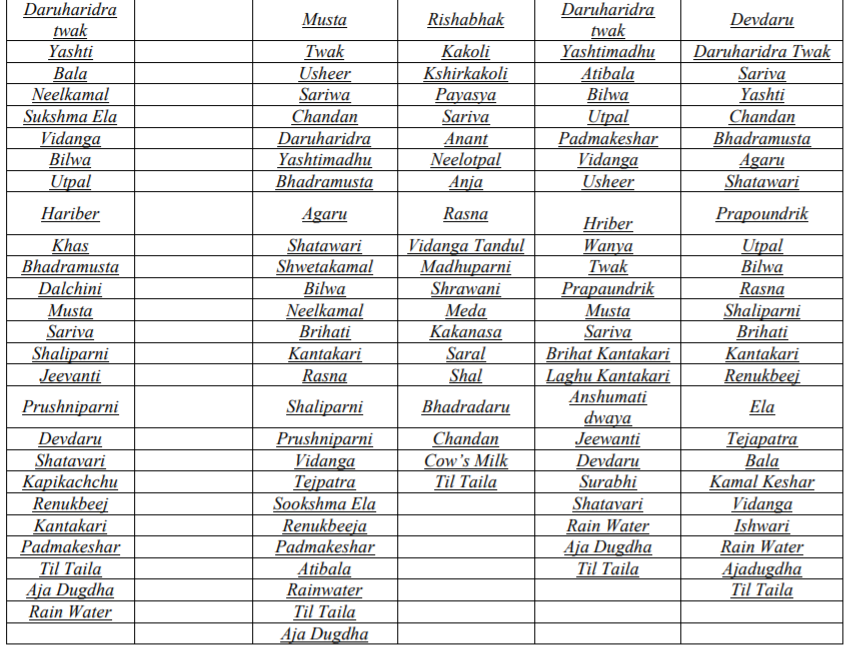

Dalvi P D, Kulkarni M S, Jadhav S P; LITERARY REVIEW OF ANU TAILA NASYA. Unique J. of Ayurvedic & Herbal Medicines. Online: www.ujconline.net Research Article ISSN 2347-2375 Received 31-01-2015; Revised 28-02-2015; Accepted 27-03-2015

It is also important to mention that in a compound Ayurvedic formulation or multi-ingredient formulation, the indications of the individual herbs have no or little role. It is the combined final effect of the formulation which is important (Ashtanga Sangraha – Sutra - 25. Chapter). Even though biological activity of the individual component of Annu oil has been mentioned below.

| **Plant component** | **Medicinal value** |
| --- | --- |
| Nāgarmothā (*Cyperus* *scariosus*) | Ayurvedic physicians use the plant for medicinal purposes for fevers, digestive system disorders, and other maladies. Modern alternative medicine recommends using the plant to treat nausea, fever and inflammation; for pain reduction; for muscle relaxation and for many other disorders.  <http://www.ijnpnd.com/article.asp?issn=2231-0738;year=2014;volume=4;issue=1;spage=23;epage=27;aulast=Imam> |
| Jīvantī (*Leptadenia* *reticulata*) | Leaves and roots of *Leptadenia* species are useful in treating skin infections and wounds. The roots are used in cardiac disease and haemorrhage, as diuretic, and to cure fever and opthalmia. They are also used as a tonic for general debility to strengthen the body. Leaves are also used as galactagogue for nursing mothers. <https://vikaspedia.in/agriculture/crop-production/package-of-practices/medicinal-and-aromatic-plants/leptadenia-reticulata> |
| Śweta candana (*Santalum* *album*) | Sandalwood oil has been widely used in folk medicine for treatment of common colds, bronchitis, skin disorders, heart ailments, general weakness, fever, infection of the urinary tract, inflammation of the mouth and pharynx, and other maladies.  Misra, Biswapriya B.; Dey, Satyahari (2013). "Evaluation of in vivoanti-hyperglycemic and antioxidant potentials of α-santalol and sandalwood oil". Phytomedicine. **20** (5): 409–16. [*doi*](https://en.wikipedia.org/wiki/Doi_(identifier)):[*10.1016/j.phymed.2012.12.017*](https://doi.org/10.1016%2Fj.phymed.2012.12.017). [*PMID*](https://en.wikipedia.org/wiki/PMID_(identifier)) [*23369343*](https://pubmed.ncbi.nlm.nih.gov/23369343).  Misra, B.B.; Dey, S. (2012). [*"Comparative phytochemical analysis and antibacterial efficacy of in vitro and in vivo extracts from East Indian sandalwood tree (Santalum album L.)"*](https://doi.org/10.1111%2Flam.12005). Letters in Applied Microbiology. **55** (6): 476–486. [*doi*](https://en.wikipedia.org/wiki/Doi_(identifier)):[*10.1111/lam.12005*](https://doi.org/10.1111%2Flam.12005). [*PMID*](https://en.wikipedia.org/wiki/PMID_(identifier)) [*23020220*](https://pubmed.ncbi.nlm.nih.gov/23020220). [*S2CID*](https://en.wikipedia.org/wiki/S2CID_(identifier)) [*36484791*](https://api.semanticscholar.org/CorpusID:36484791). |
| Jala (*Pavonia* *odorata*) | *Pavonia odorata* is a aromatic medicinal herb used in the treatment of fever, burning sensation, herpes, excessive thirst, giddiness, vitiligo, vomiting, diarrhea, etc. Its fragrant root is used as hair tonic.  <https://www.easyayurveda.com/2019/05/04/hrivera-pavonia-odorata> |
| Pṛśniparṇī (*Uraria* *picta*) | It is used as a cardio and nervine tonic and has anti-inflammatory, expectorant, and diuretic properties. The root of the plant is one of the ingredients of ‘dasamoola’ in Ayurveda. <https://vikaspedia.in/agriculture/crop-production/package-of-practices/medicinal-and-aromatic-plants/uraria-picta> |
| Bela (*Aegle* *marmelos*) | The leaves, bark, roots, fruits, and seeds are used in traditional medicine to treat various illnesses  Misra KK (1999). ["Bael"](http://www.hort.purdue.edu/newcrop/CropFactSheets/bael.html). NewCROP, the New Crop Resource Online Program, Department of Horticulture and Landscape Architecture, Center for New Crops & Plant Products, Purdue University, W. Lafayette, IN. Retrieved 20 January 2016 |
| Devdāru (*Cedrus* *deodara*) | The Deodar Cedar has many medicinal properties. Among other things, it is stated that:The extract of essential oil from tree wood has a significant  **anti-inflammatory** effect. The Cedrus deodara branch’s bark air-dried aqueous extract has an **antiarthritic** activity <https://www.kalliergeia.com/en/deodar-cedar-cedrus-deodara-description-and-uses/> |
| Dāruharidrā (*Berberis* *aristata*) | In India, *B. aristata* is used in traditional herbal medicine. Its stem, roots, and fruits are used in [Ayurveda](https://en.wikipedia.org/wiki/Ayurveda). The root bark contains the bitter alkaloid [berberine](https://en.wikipedia.org/wiki/Berberine), which has been studied for its potential pharmacological properties.  Kala, C.P.; et al. (2006). [*"Developing the medicinal plants sector in northern India: challenges and opportunities"*](https://www.ncbi.nlm.nih.gov/pmc/articles/PMC1562365). Journal of Ethnobiology and Ethnomedicine. **2**: 32. [*doi*](https://en.wikipedia.org/wiki/Doi_(identifier)):[*10.1186/1746-4269-2-32*](https://doi.org/10.1186%2F1746-4269-2-32). [*PMC*](https://en.wikipedia.org/wiki/PMC_(identifier)) [*1562365*](https://www.ncbi.nlm.nih.gov/pmc/articles/PMC1562365).  Parmar, C. & M.K. Kaushal (1982). [*Berberis aristata*](http://www.hort.purdue.edu/newcrop/parmar/03.html). *Wild Fruits*. New Delhi, India: Kalyani Publishers. pp. 10–14. |
| Tejpatra (*Cinnamomum* *tamala*) | Its leaves have a clove-like aroma with a hint of peppery taste; they are used for culinary and medicinal purposes. |
| Kamala keṣara (*Nelumbo* *nucifera*) | All parts of *Nelumbo nucifera* are edible, with the rhizome and seeds being the main consumption parts. Traditionally rhizomes, leaves, and seeds have been used as [folk medicines](https://en.wikipedia.org/wiki/Traditional_medicine), [Ayurveda](https://en.wikipedia.org/wiki/Ayurveda), [Chinese traditional medicine](https://en.wikipedia.org/wiki/Traditional_Chinese_medicine), and [oriental medicine](https://en.wikipedia.org/wiki/Oriental_medicine). While leaves are used for [hematemesis](https://en.wikipedia.org/wiki/Hematemesis), [epistaxis](https://en.wikipedia.org/wiki/Nosebleed), and [hematuria](https://en.wikipedia.org/wiki/Hematuria" \o "Hematuria), the flowers are used for lowering blood sugar levels, [diarrhea](https://en.wikipedia.org/wiki/Diarrhea" \o "Diarrhea), [cholera](https://en.wikipedia.org/wiki/Cholera), [fever](https://en.wikipedia.org/wiki/Fever), and hyperdipsia. Rhizomes are promoted have purported [diuretic](https://en.wikipedia.org/wiki/Diuretic), [antidiabetic](https://en.wikipedia.org/wiki/Anti-diabetic_medication), and [anti-inflammatory](https://en.wikipedia.org/wiki/Anti-inflammatory) properties.  Khare CP. *Indian Herbal Remedies: Rational Western Therapy, Ayurvedic, and Other Traditional Usage, Botany*, 1st edn. USA: Springer, 2004: 326–327.  KR, Bhat R. Lotus: a potential nutraceutical source. *J Agri Technol* 2007; **3**: 143–155 |
| Sevya (*Chrysopogon* *zizanioides*) | It is used for its antiseptic properties to treat acne and sores  [Technical Bulletin No. 2001/1"](https://web.archive.org/web/20130918112455/http:/www.vetiver.com/PRVN_med_aro%20doc.pdf). Pacific Rim Vetiver Network. September 2001. Archived from [the original](http://www.vetiver.com/PRVN_med_aro%20doc.pdf) on 2013-09-18. Retrieved 2008-07-07 |
| Viḍañga (*Embelia* *ribes*) | In [Ayurveda](https://en.wikipedia.org/wiki/Ayurveda) and [Siddha](https://en.wikipedia.org/wiki/Siddha), it is considered widely beneficial in variety of diseases In particular [embelin](https://en.wikipedia.org/wiki/Embelin) isolated from dried berries of Embelia ribes has a wide spectrum of biological activities. |
| Utpala (*Nymphaeanouchali*) | *N. nouchali* is considered a medicinal plant in Indian Ayurvedic medicine under the name *ambal*; it was mainly used to treat indigestion.  P. V. Sharma, *Puṣpāyurvedaḥ* - Pradhāna vitaraka Caukhambhā Bhāratī Akādamī, 1998 |
| Anantmūla (*Hemidesmus* *indicus*) | The extracts from the root are used in syrup with sugar and a dash of lemon (Sharbat) and served at most small refreshment shops in South India. |
| Tila Taila (*Sesamum indicum*) | A meta-analysis showed that sesame consumption produced small reductions in both systolic and diastolic blood pressure. Sesame oil studies reported a reduction of oxidative stress markers and lipid peroxidation.  Khosravi-Boroujeni H, Nikbakht E, Natanelov E, Khalesi S (2017). "Can sesame consumption improve blood pressure? A systematic review and meta-analysis of controlled trials". [*Journal of the Science of Food and Agriculture*](https://en.wikipedia.org/wiki/Journal_of_the_Science_of_Food_and_Agriculture). **97** (10): 3087–3094. [*doi*](https://en.wikipedia.org/wiki/Doi_(identifier)):[*10.1002/jsfa.8361*](https://doi.org/10.1002%2Fjsfa.8361). [*PMID*](https://en.wikipedia.org/wiki/PMID_(identifier)) [*28387047*](https://pubmed.ncbi.nlm.nih.gov/28387047).  Gouveia Lde A, Cardoso CA, de Oliveira GM, Rosa G, Moreira AS (2016). "Effects of the Intake of Sesame Seeds (Sesamum indicumL.) and Derivatives on Oxidative Stress: A Systematic Review". Journal of Medicinal Food. **19** (4): 337–345. [*doi*](https://en.wikipedia.org/wiki/Doi_(identifier)):[*10.1089/jmf.2015.0075*](https://doi.org/10.1089%2Fjmf.2015.0075). [*PMID*](https://en.wikipedia.org/wiki/PMID_(identifier)) [*27074618*](https://pubmed.ncbi.nlm.nih.gov/27074618) |
| Muleṭhī (*Glycyrrhiza* *glabra*) | In [traditional Chinese medicine](https://en.wikipedia.org/wiki/Traditional_Chinese_medicine), a related species [*G. uralensis*](https://en.wikipedia.org/wiki/Glycyrrhiza_uralensis) (often translated as "liquorice") is known as "gancao" ([Chinese](https://en.wikipedia.org/wiki/Chinese_language): 甘草; "sweet grass"), and is believed to "harmonize" the ingredients in a formula.  Bensky, Dan; et al. (2004). Chinese Herbal Medicine: Materia Medica, Third Edition. Eastland Press. [*ISBN*](https://en.wikipedia.org/wiki/ISBN_(identifier)) [*978-0-939616-42-8*](https://en.wikipedia.org/wiki/Special:BookSources/978-0-939616-42-8).  Balakrishna, Acharya (2006). Ayurveda: Its Principles & Philosophies. New Delhi, India: Divya prakashan. p. 206. [*ISBN*](https://en.wikipedia.org/wiki/ISBN_(identifier)) [*978-8189235567*](https://en.wikipedia.org/wiki/Special:BookSources/978-8189235567).  Tewari, D; Mocan, A; Parvanov, E. D; Sah, A. N; Nabavi, S. M; Huminiecki, L; Ma, Z. F; Lee, Y. Y; Horbańczuk, J. O; Atanasov, A. G (2017). [*"Ethnopharmacological Approaches for Therapy of Jaundice: Part II. Highly Used Plant Species from Acanthaceae, Euphorbiaceae, Asteraceae, Combretaceae, and Fabaceae Families"*](https://www.ncbi.nlm.nih.gov/pmc/articles/PMC5554347). Frontiers in Pharmacology. **8**: 519. [*doi*](https://en.wikipedia.org/wiki/Doi_(identifier)):[*10.3389/fphar.2017.00519*](https://doi.org/10.3389%2Ffphar.2017.00519). [*PMC*](https://en.wikipedia.org/wiki/PMC_(identifier)) [*5554347*](https://www.ncbi.nlm.nih.gov/pmc/articles/PMC5554347). [*PMID*](https://en.wikipedia.org/wiki/PMID_(identifier)) [*28848436*](https://pubmed.ncbi.nlm.nih.gov/28848436).  Wendy Christensen (2009). [*Empire of Ancient Egypt*](https://books.google.com/books?id=LpQBdsH12acC&pg=PA98). Infobase Publishing. pp. 98–. [*ISBN*](https://en.wikipedia.org/wiki/ISBN_(identifier)) [*978-1-60413-160-4*](https://en.wikipedia.org/wiki/Special:BookSources/978-1-60413-160-4). |
| Śatāvarī (*Asparagus* *racemosus*) | Shatavari is important in traditional [Ayurvedic medicine](https://en.wikipedia.org/wiki/Ayurveda). The key pharmacologic constituents of shatavari are steroidal [saponins](https://en.wikipedia.org/wiki/Saponin" \o "Saponin), [mucilage](https://en.wikipedia.org/wiki/Mucilage), and [alkaloids](https://en.wikipedia.org/wiki/Alkaloid).  Pizzorno Jr., Joseph E.; Murray, Michael T.; Joiner-Bey, Herb (2015). The Clinician's Handbook of Natural Medicine (3rd ed.). Churchill Livingstone. p. 516. [*ISBN*](https://en.wikipedia.org/wiki/ISBN_(identifier)) [*9780702055140*](https://en.wikipedia.org/wiki/Special:BookSources/9780702055140).  Hechtman, Leah (2018). Clinical Naturopathic Medicine (2 ed.). Elsevier. pp. 879, 908. [*ISBN*](https://en.wikipedia.org/wiki/ISBN_(identifier)) [*9780729542425*](https://en.wikipedia.org/wiki/Special:BookSources/9780729542425).  Goyal, R. K.; Singh, Janardhan; Lal, Harbans (September 2003). "Asparagus racemosus—an update". Indian Journal of Medical Sciences. **57** (9): 408–414. [*PMID*](https://en.wikipedia.org/wiki/PMID_(identifier)) [*14515032*](https://pubmed.ncbi.nlm.nih.gov/14515032).  [*"Wild Asparagus"*](https://toxnet.nlm.nih.gov/cgi-bin/sis/search2/f?./temp/~R2Mxse:1). LactMed. National Institute of Health*. Retrieved 13 November 2018*. |
| Bṛhatī (*Solanum* *indicum*) | The Solanum plant is reported to be bitter, acrid, astringent, carminative, stomachic, resolvent, demulcent, diuretic, emmenagogue, febrifuge, and cardiotonic. It is useful in the treatment of asthma, catarrh, dropsy, chest pain, chronic fever, colic, dry and spasmodic cough, <https://vikaspedia.in/agriculture/crop-production/package-of-practices/medicinal-and-aromatic-plants/solanum-indicum> |
| Kaṇṭakārī (*Solanum* *surattense*) | Panchang (whole herb including roots) and berries, have anthelmintic property, useful in bronchitis, asthma, fever relieving, thirst and given in urinary concretions. <https://vikaspedia.in/agriculture/crop-production/package-of-practices/medicinal-and-aromatic-plants/solanum-surattense-1> |
| Surbhi (*Pluchea* *lanceolata*), | Whole plant is used in Ayurvedic medicine. *Pluchea lanceolata* is accepted as classical drug for arthritis. Its decoction is given for rheumatic conditions, muscular pains, edema, and fever and also applied externally as massage oil. <https://vikaspedia.in/agriculture/crop-production/package-of-practices/medicinal-and-aromatic-plants/pluchea-lanceolata> |
| Śālaparṇī (*Desmodium* *gangeticum*), | Medically, the plant has many benefits. The plant is deemed to restore proper functioning of the body by increasing health and vitality, supporting the structure of organ tissue, reduce fever and cough, and support digestion. "*Pleurolobus gangeticus*". [*Germplasm Resources Information Network*](https://en.wikipedia.org/wiki/Germplasm_Resources_Information_Network) *(GRIN)*. [Agricultural Research Service](https://en.wikipedia.org/wiki/Agricultural_Research_Service) (ARS), [United States Department of Agriculture](https://en.wikipedia.org/wiki/United_States_Department_of_Agriculture) (USDA) |
| Truṭi (*Elettaria* cardamomum), | True cardamom may have been used in [Ayurveda](https://en.wikipedia.org/wiki/Ayurveda) medicine as early as the 4th century BC.  [Pickersgill, Barbara](https://en.wikipedia.org/wiki/Barbara_Pickersgill) (2005). Prance, Ghillean; Nesbitt, Mark (eds.). *The Cultural History of Plants*. Routledge. p. 158. [ISBN](https://en.wikipedia.org/wiki/ISBN_(identifier)) [0415927463](https://en.wikipedia.org/wiki/Special:BookSources/0415927463). |
| Reṇukā (*Vitex agnus-castus*), | Verkaik S, Kamperman AM, van Westrhenen R, Schulte PF (2017). "The treatment of premenstrual syndrome with preparations of Vitex agnus castus: a systematic review and meta-analysis". American Journal of Obstetrics and Gynecology. **217** (2): 150166. [*doi*](https://en.wikipedia.org/wiki/Doi_(identifier)):[*10.1016/j.ajog.2017.02.028*](https://doi.org/10.1016%2Fj.ajog.2017.02.028). [*PMID*](https://en.wikipedia.org/wiki/PMID_(identifier)) [*28237870*](https://pubmed.ncbi.nlm.nih.gov/28237870). [*S2CID*](https://en.wikipedia.org/wiki/S2CID_(identifier)) [*41615874*](https://api.semanticscholar.org/CorpusID:41615874).  Van Die, M. D; Burger, H. G; Teede, H. J; Bone, K. M (2013). [*"Vitex agnus-castus extracts for female reproductive disorders: A systematic review of clinical trials"*](https://doi.org/10.1055%2Fs-0032-1327831). Planta Medica. **79** (7): 562–75. [*doi*](https://en.wikipedia.org/wiki/Doi_(identifier)):[*10.1055/s-0032-1327831*](https://doi.org/10.1055%2Fs-0032-1327831). [*PMID*](https://en.wikipedia.org/wiki/PMID_(identifier)) [*23136064*](https://pubmed.ncbi.nlm.nih.gov/23136064).  Daniele, C; Thompson Coon, J; Pittler, M. H.; Ernst, E (2005). "Vitex agnus castus: a systematic review of adverse events". Drug Safety. **28** (4): 319–32. [*doi*](https://en.wikipedia.org/wiki/Doi_(identifier)):[*10.2165/00002018-200528040-00004*](https://doi.org/10.2165%2F00002018-200528040-00004). [*PMID*](https://en.wikipedia.org/wiki/PMID_(identifier)) [*15783241*](https://pubmed.ncbi.nlm.nih.gov/15783241). [*S2CID*](https://en.wikipedia.org/wiki/S2CID_(identifier)) [*45851264*](https://api.semanticscholar.org/CorpusID:45851264)  ["Chaste tree"](https://www.drugs.com/npp/chaste-tree.html). Drugs.com. 9 October 2017. Retrieved 20 August 2019. |
| Ajadugdha (Goat milk) | It heals and rejuvenates the body, treats respiratory diseases, has antibacterial and bactericidal effects. It improves the activity of the cardiovascular system. It strengthens the immune system.  <https://mahabazar.club/en/aja-dugdha/> |
